# Method for cycle detection in sparse, irregularly sampled, long-term neuro-behavioral timeseries: Basis pursuit denoising with polynomial detrending of long-term, inter-ictal epileptiform activity

**DOI:** 10.1101/2023.05.04.539355

**Authors:** Irena Balzekas, Joshua Trzasko, Grace Yu, Thomas J. Richner, Filip Mivalt, Vladimir Sladky, Nicholas M. Gregg, Jamie Van Gompel, Kai Miller, Paul E. Croarkin, Vaclav Kremen, Gregory A. Worrell

**Affiliations:** Bioelectronics, Neurophysiology, and Engineering Laboratory, Department of Neurology, Mayo Clinic, Rochester, MN, United States; Biomedical Engineering and Physiology Graduate Program, Mayo Clinic Graduate School of Biomedical Sciences, Rochester, MN, United States; Mayo Clinic Alix School of Medicine, Rochester, MN, United States; Mayo Clinic Medical Scientist Training Program, Rochester, MN, United States; Faculty of Biomedical Engineering, Czech Technical University in Prague, Kladno, Czechia; Faculty of Electrical Engineering and Communication, Department of Biomedical Engineering, Brno University of Technology, Brno, Czechia; Department of Psychiatry and Psychology, Mayo Clinic, Rochester, MN, United States; Department of Neurosurgery, Mayo Clinic, Rochester, MN, United States; Czech Institute of Informatics, Robotics and Cybernetics, Czech Technical University in Prague, Prague, Czechia; Department of Radiology, Mayo Clinic, Rochester, MN, United States; International Clinic Research Center, St. Anne’s University Research Hospital, Brno, Czech Republic; Faculty of Biomedical Engineering, Czech Technical University in Prague, Prague, Czech Republic

## Abstract

Numerous physiological processes are cyclical, but sampling these processes densely enough to perform frequency decomposition and subsequent analyses can be challenging. Mathematical approaches for decomposition and reconstruction of sparsely and irregularly sampled signals are well established but have been under-utilized in physiological applications. We developed a basis pursuit denoising with polynomial detrending (BPWP) model that recovers oscillations and trends from sparse and irregularly sampled timeseries. We validated this model on a unique dataset of long-term inter-ictal epileptiform discharge (IED) rates from human hippocampus recorded with a novel investigational device with continuous local field potential sensing. IED rates have well established circadian and multiday cycles related to sleep, wakefulness, and seizure clusters. Given sparse and irregular samples of IED rates from multi-month intracranial EEG recordings from ambulatory humans, we used BPWP to compute narrowband spectral power and polynomial trend coefficients and identify IED rate cycles in three subjects. In select cases, we propose that random and irregular sampling may be leveraged for frequency decomposition of physiological signals.

**Author Summary:** Circadian and multiday cycles are an important part of many long-term neuro-behavioral phenomena such as pathological inter-ictal epileptiform discharges (IEDs) in epilepsy. Long-term, ambulatory, neuro-behavioral monitoring in human patients involves complex recording systems that can be subject to intermittent, irregular data loss and storage limitations, resulting in sparse, irregularly sampled data. Cycle identification in sparse data or irregular data using traditional frequency decomposition techniques typically requires interpolation to create a regular timeseries. Using unique, long-term recordings of pathological brain activity in patients with epilepsy implanted with an investigational device, we developed a method to identify cycles in sparse, irregular neuro-behavioral data without interpolation. We anticipate this approach will enable retrospective cycle identification in sparse neuro-behavioral timeseries and support prospective sparse sampling in monitoring systems to enable long-term monitoring of patients and to extend storage capacity in a variety of ambulatory monitoring applications.

## Introduction

Studying the dynamics of complex neurological and psychiatric diseases with behavioral manifestations requires ambulatory monitoring of peripheral physiology and brain activity. In epilepsy, long-term monitoring of seizures and inter-ictal epileptiform discharges (IEDs) has revealed circadian and multiday cycles of seizures and epileptiform brain activity[1-3]. In temporal lobe epilepsy (TLE) seizures and IED rates are especially tied to sleep/wake behavioral states[3-7]. Leveraging these cycles to anticipate high risk intervals for seizure occurrence and timw interventions may create new therapeutic opportunities[3].

Data under-sampling and loss are common in ambulatory monitoring, especially for devices recording intracranial electroencephalography (iEEG). Complex recording systems (Fig. 1a) are subject to signal storage, battery life, and wireless connectivity limitations[8]. Given device data storage limitations, most electrical brain stimulation (EBS) devices in clinical use store variable and limited local field potential (LFP) data: around one month of coarsely averaged features such as power-in-band, for example[9, 10] or limited brief selected raw data segments [9, 10]. Multiday drops tend to present the largest analytical challenges. Often, past data are overwritten if recordings are not downloaded by patients[9]. Research grade, investigational devices capable of continuous data streaming require frequent charging and consistent wireless connectivity; creating significant hardware management burdens for patients[8, 11, 12]. Power and connectivity issues lead to packet drops and data loss that can degrade analytics and clinical performance[13]. Approaches that maximize the collection of meaningful data while minimizing demands on patients and system hardware are needed.

**Figure 1.**
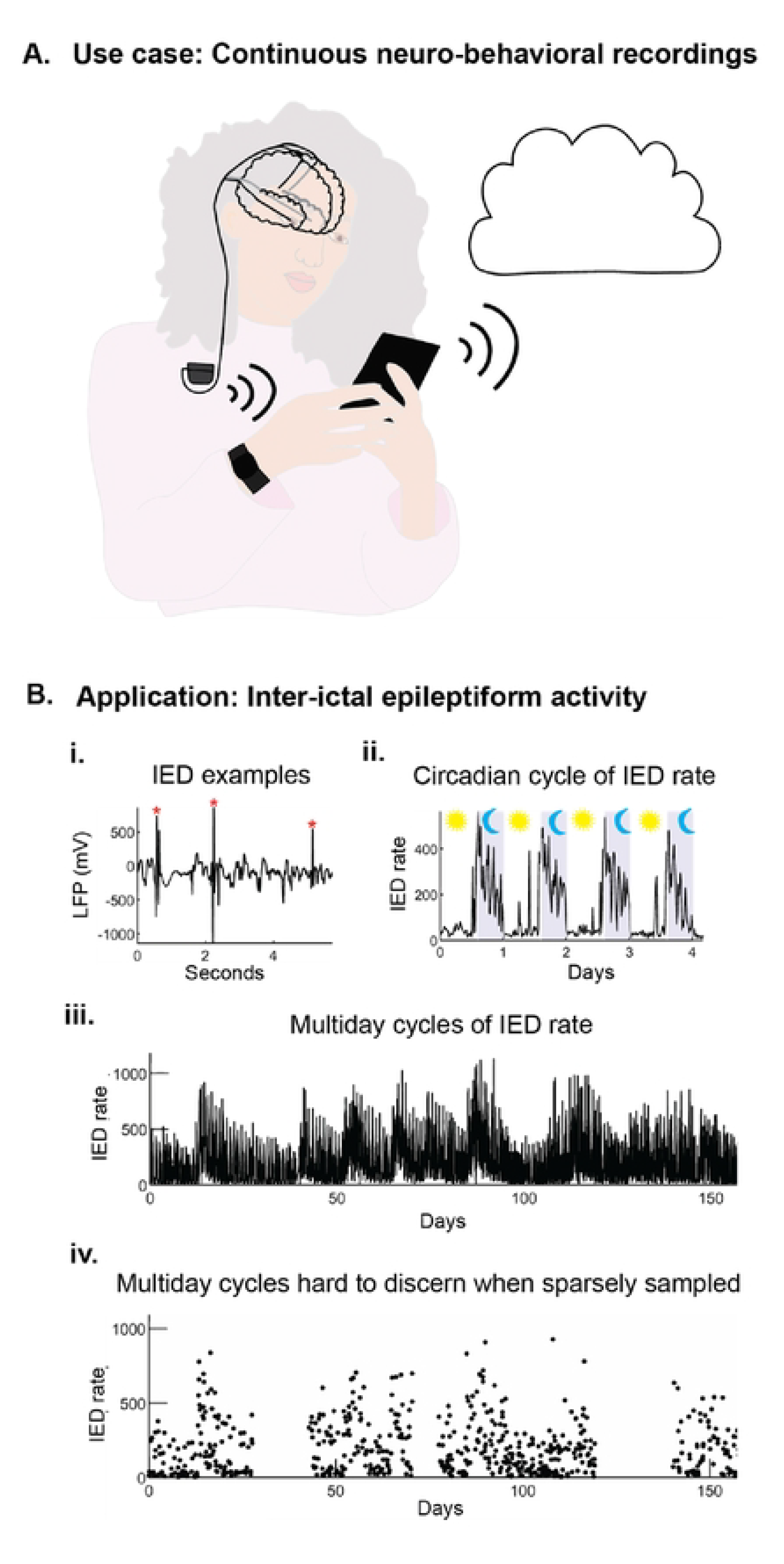
Schematic overview of neuro-behavioral recording applications. A) Use case: Systerns for continuous neuro-behavioral recordings. Recording, energy, storage, connectivity, and usability dernands placed on arnbulatory brain recording systen1s can create data loss. B) Application: Cyclical neuro-behavioral signals - lnter-ictal epileptiform activity. i) Example of inter-ictal epileptifonn discharges (IEDs) recorded frorn intracranial electroencephalography (iEEG). ii) Exarnple ofIED rates (IED per hour) recorded frorn a hippocarnpal iEEG electrode showing circadian changes in IED rate with increased lED rates during sleep. The period from IOPM to 8 AM is noted in gray. iii) Exarnple of multiday IED rate recording shov,ing 1nultiday cycles. iv) Randon1 subsan1pling of the signal in iii to show how sparse sarnpling and data drops rnake it challenging to discern the underlying cycles of IED rate.

**Figure 2:**
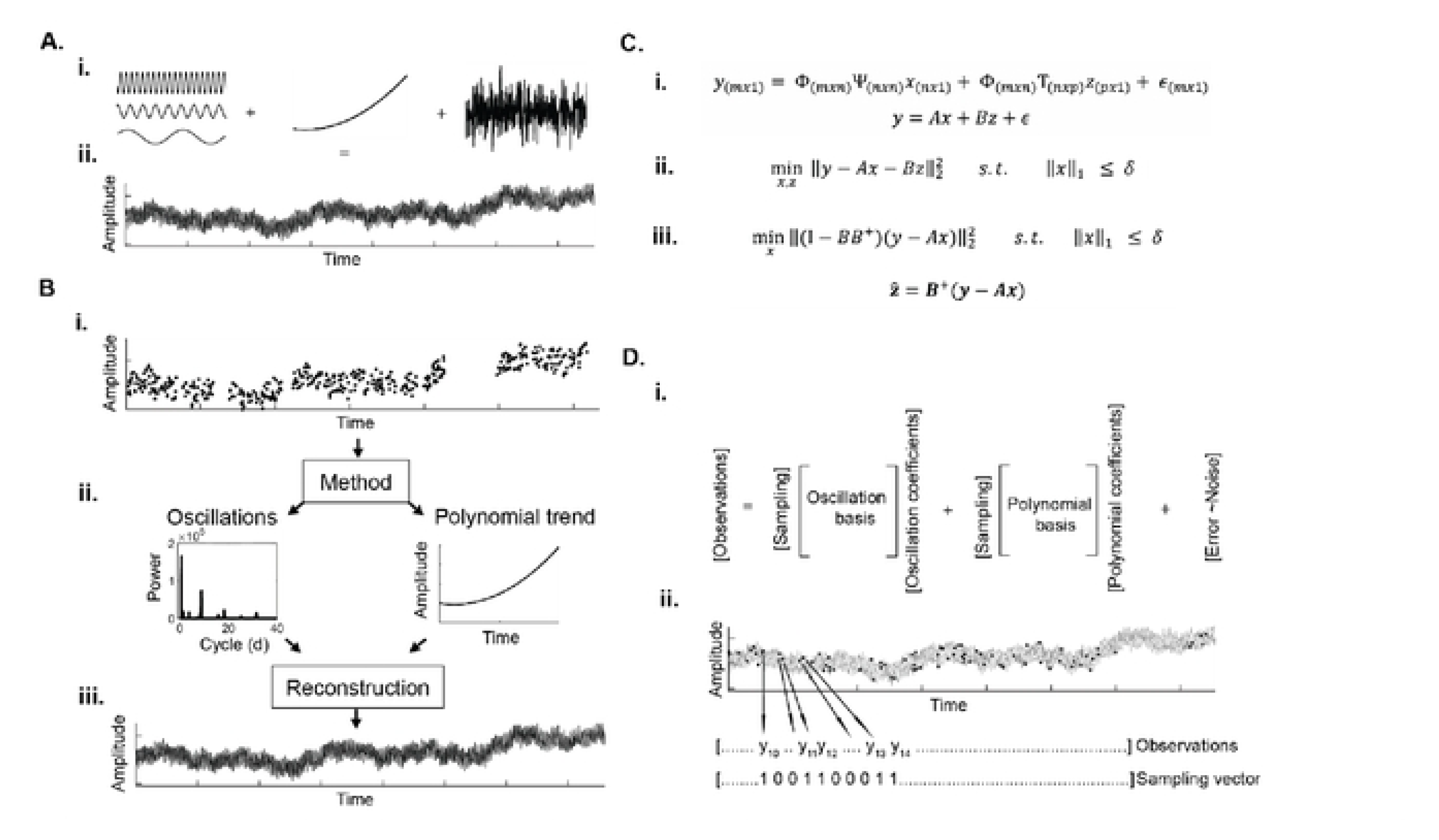
Description of method objectives and signal assurnptions. A) The core assurnption of the rnodel is that the underlying signal (ii) is the linear surn of (i) oscillations, a polynornial trend, and noise. Part B) describes the overall workflow including (i) data input to the model, (ii) outputs, and (iii) estirnated signal reconstructions. Ci) Equation representing the core signal assurnption that the observations come from a combination of oscillations, a polynomial trend, and noise or error. Notation includes *y* (rn x I vector of observed data), Ψ, (n x n discrete cosine transfonn (OCT) basis), Φ (m x n binary row subsampling matrix), *x* (n x I OCT coefficients), T (n x p Vandermonde n1atrix), *z* (p x I polynomial coefficients), ε (rn x I error tenns). Cii) Expression for basis pursuit denoising containing *x* and *z* as unknowns, yielding a 20 1nini111ization problern that is reduced to a ID minin1ization problern (iii) by variable projection. D) Schematic representation of the equations and sampling approach in C.

One solution is to store only the minimum data required to characterize a neurophysiological process[14]. An acceptable, minimal representation of data depends on the signal of interest and requirements of methods for analysis[15]. Most approaches for cycle identification are unsuitable for infrequently and irregularly sampled timeseries. Classical Fourier-[16] and autocorrelation-based[17] methods require dense and regular sampling, increasing storage demands and necessitating interpolation of missing data. Identifying cycles in timeseries wherein most samples are unknown requires approaches designed to accommodate sparsity.

Sparsity can describe datasets, signals, and statistical approximations. These definitions are often concurrently relevant. In a sparse dataset, most observations, or samples, are zero. Sometimes, a signal can be well represented by a small number of components (low rank approximation) that capture nearly all the variance of a signal. In this case the signal, and its model, are both sparse. There are mathematical definitions whereby values describe the number of nonzero coefficients. For example, a signal is k-sparse if only k coefficients are nonzero. In sparse representations, the value of k is small, though its exact value is case dependent. This can also be explained as sparsity in a transform domain. For example, although a signal may have been sampled at a high sampling rate to create a very dense dataset in the time domain, in the frequency domain, the power spectrum may only have 3, narrow peaks indicating that what was observed was 3, separable, concurrent oscillations. Altogether, a signal that has a few, distinctive, separable components that capture most of the signal’s variance lends itself well to sparse representation and ultimately analysis or reconstruction of a sparse dataset using sparsity- (or low-rank)-based techniques.

Sparse recovery techniques (including compressed sensing methods) capture a family of techniques developed to recover information from sparse and sparsely sampled data in applications including signal compression and magnetic resonance imaging (MRI)[18-20]. Based on basis pursuit[21], these methods exploit conditions whereby algebraically undetermined or sub-Nyquist-Shannon sampling is permissible and frequency decomposition can be done on less frequently sampled signals[19]. The related techniques of basis pursuit denoising and least absolute shrinkage selection operator (LASSO), also known as L1 regularization or sparse regression, estimate the simplest, or in this case the sparsest, description for a set of observations that does not have to be regularly or densely sampled[22, 23]. By leveraging sparsity of the target signal quantity, we can address the challenge presented by long-term neuro-behavioral data acquisition: identifying cycles in infrequently and irregularly sampled data[24]. Our goal was to develop a conservative model for frequency decomposition of infrequently and irregularly sampled data that yields sparse spectral outputs and accommodates polynomial trends in the timeseries, limiting potential errors introduced by linear detrending during pre-processing.

Here, we describe a method for frequency decomposition of sparsely and irregularly sampled neuro-behavioral data: basis pursuit denoising with polynomial detrending (BPWP). In short, the approach takes an irregularly and sparsely sampled timeseries and performs frequency decomposition and polynomial detrending. We used simulations to test the method’s characterization of cycles in complex timeseries data over a range of physiologically relevant parameters. We then applied BPWP to unique, real-world recordings of chronic, ambulatory, iEEG from patients implanted with the investigational Medtronic Summit RC+S™ deep brain stimulation (DBS) device for drug resistant temporal lobe epilepsy (TLE). The recordings contained multiple years of translationally relevant IED rates from epileptic hippocampus in 3 human participants. We identified circadian and multiday cycles of IED rate in our data with both standard approaches and BPWP. Our findings support frequency decomposition of sparsely sampled neuro-behavioral data and highlight translational opportunities for efficient sampling and reconstruction of neuro-behavioral signals to study the interplay between electrophysiology and behavioral or multi-domain data.

## Results

The purpose of BPWP is to identify oscillations and polynomial trends in sparsely and irregularly sampled timeseries. Explained in detail in the methods, the model assumes the observed signal (a set of sparse, irregular samples) can be described by the linear sum of a set of oscillations, a polynomial trend, and noise. BPWP aims to minimize the square of the L2 norm of the difference between the observed data and the estimate of the observed data (the sum of the oscillations, polynomial trend, and noise). The model outputs consist of a sparse, narrowband power spectrum (spectral coefficients mapping to discrete cosine transform (DCT) basis) and a polynomial trend. BPWP constrains the number and amplitude of the spectral coefficients by restricting the L1 norm of the coefficient vector to be less than the parameter *δ*. The *δ* parameter is selected in a subject-specific manner via parameter sweeps to identify the *δ* value that minimizes the mean square error between the observed data and the model estimates. The spectrum and polynomial trend are then used to estimate the regular, dense representation of the signal that yielded the sparse, irregular samples. We first demonstrate the method on a simulated IED-rate timeseries with pre-defined spectral composition. We then test the method on real-world IED rate data by taking the densely sampled long-term, real-world timeseries, sparsely sampling it at different densities and paradigms, and applying BPWP.

### Cycle identification in simulated IED-rate timeseries

The simulated IED rate timeseries is shown in Figure 3a. The CWT spectrogram (Fig. 3b) and average CWT spectrum (Fig. 3f) reflect the increased signal power at periods of one day and one month. The results of the 10-fold 75/25 cross-validation (Fig. 3f) for *δ* parameter selection capture a clear optimal *δ* value that minimizes mean square error (MSE) for each random sampling rate tested.

**Figure 3.**
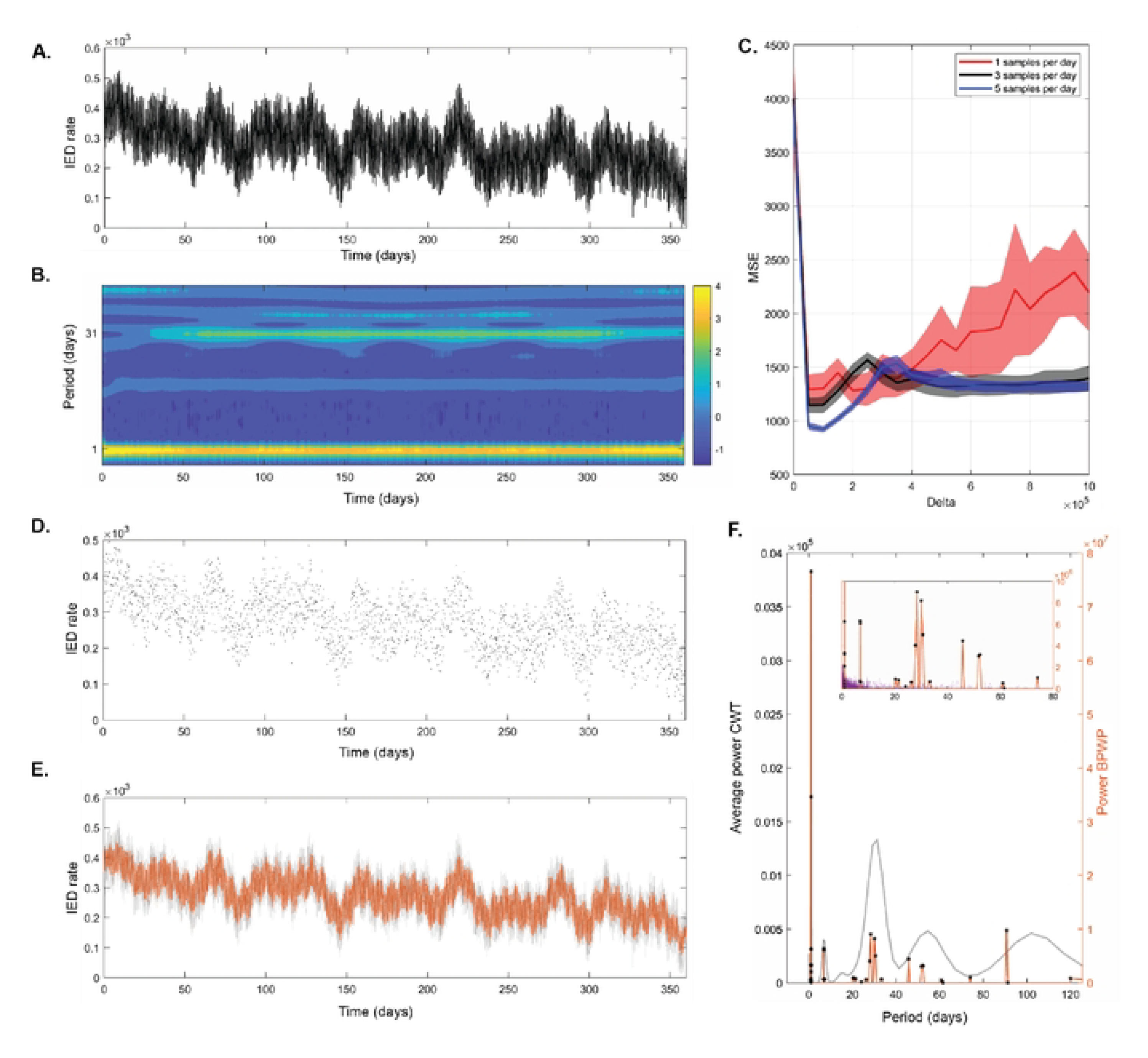
BPWP outputs for sirnulated JED timeseries. A) Raw data show hourly rate ofIEDs, updated every 20 rninutes, from a simulated signal containing oscillations at I, 7, IS, 21, 30, SO, and I00 days. Tin1eseries consists of over 20,000 samples. B) Cornplex wavelet transform (CWT) spectrogran1 of the tirneseries in A showing power in different cycles. Colorbar indicates spectral power. C) Results often-fold 75/25 cross-validation for *(5* parameter selection. Average rnean square error with 95% confidence intervals for each *(5* value and sarnpling rate tested are shown. D) Randon1 sarnples frorn the raw data in A averaging at five sarnples per day. Timeseries consists of 1,800 total samples. E) Underlying raw data are shown in gray. The estirnated reconstruction of the underlying data based on the sparse samples in Dis shown in orange. **F)** The average CWT spectnnn (average over tirne frorn the spectrogran1 in B) is shown in gray. The method’s spectral output is shown in orange. Black stars denote significant peaks; peaks whose arnplitude was above the 99^th^ percentile of the distribution created by shuffling the input data and re-calculating the method I00 tin1es. The insert shows the spectral outputs from the reshuffling in purple. The narrowband peaks from the method align with the central tendencies of the broadband CWT peaks.

The BPWP power spectrum (Fig. 3f) derived from 5 random samples per day (1,800 total samples) (Fig. 3d) shows significant peaks aligned with the CWT spectrum (derived from >26,000 samples). The insert in Figure 3f shows significant BPWP peaks in relation to the noise floor (purple) which was determined by reshuffling the samples and recalculating BPWP. The amplitude of the noise floor is nearly an order of magnitude lower than the significant BPWP outputs. The signal reconstruction based on BPWP spectral and polynomial coefficient estimates (Fig. 3e) closely captures both the multiday cycles and polynomial trend present in the simulated timeseries.

The reliability of cycle detection by BPWP depends on cycle length and on the variance and signal to noise ratio (SNR) of the underlying signal. We simulated oscillations with cycle lengths from 1 to 120 days with high and low variance and SNR conditions (Supplementary methods, Supplementary fig. 2ai, bi). Repeat sampling and recalculation of BPWP of these oscillations were used to determine the percent detection rate of the known oscillation in each condition. The reliability of cycle detection was worst in the low variance, low SNR, and lowest sampling rate conditions (Supplementary fig. 2). At a low sampling rate, a relationship between cycle length and detection rate becomes evident; detection rate decreases with increasing cycle period (Supplementary fig. 2).

### Cycle identification in real-world IED rate timeseries

The results of 10-fold 75/25 cross validation to select a *δ* parameter for each subject are shown in Supplementary Figure 1. The values of *δ* that yielded the lowest MSE for different sampling rates were consistent within subjects. Although the optimal *δ* value was in a similar range between subjects, we selected patient-specific *δ* values to minimize under or over-fitting by fixing *δ* across subjects.

The raw, hourly IED rate data from participant 1 show multiday cycles and a gradual increase in rate over time (Fig. 4a). This is reflected in the CWT spectrogram (Fig. 4b) of the IED timeseries which illustrates the stable circadian cycle and multiday cycles around two and four weeks. The average CWT spectrum across time is depicted in Figure 4c and demonstrates that this participant’s IED rate component oscillations include periods around one, 18, 30, 50, and 100 days. Raw IED rate timeseries for the other participants are available in supplementary figures 3 and 4.

**Figure 4:**
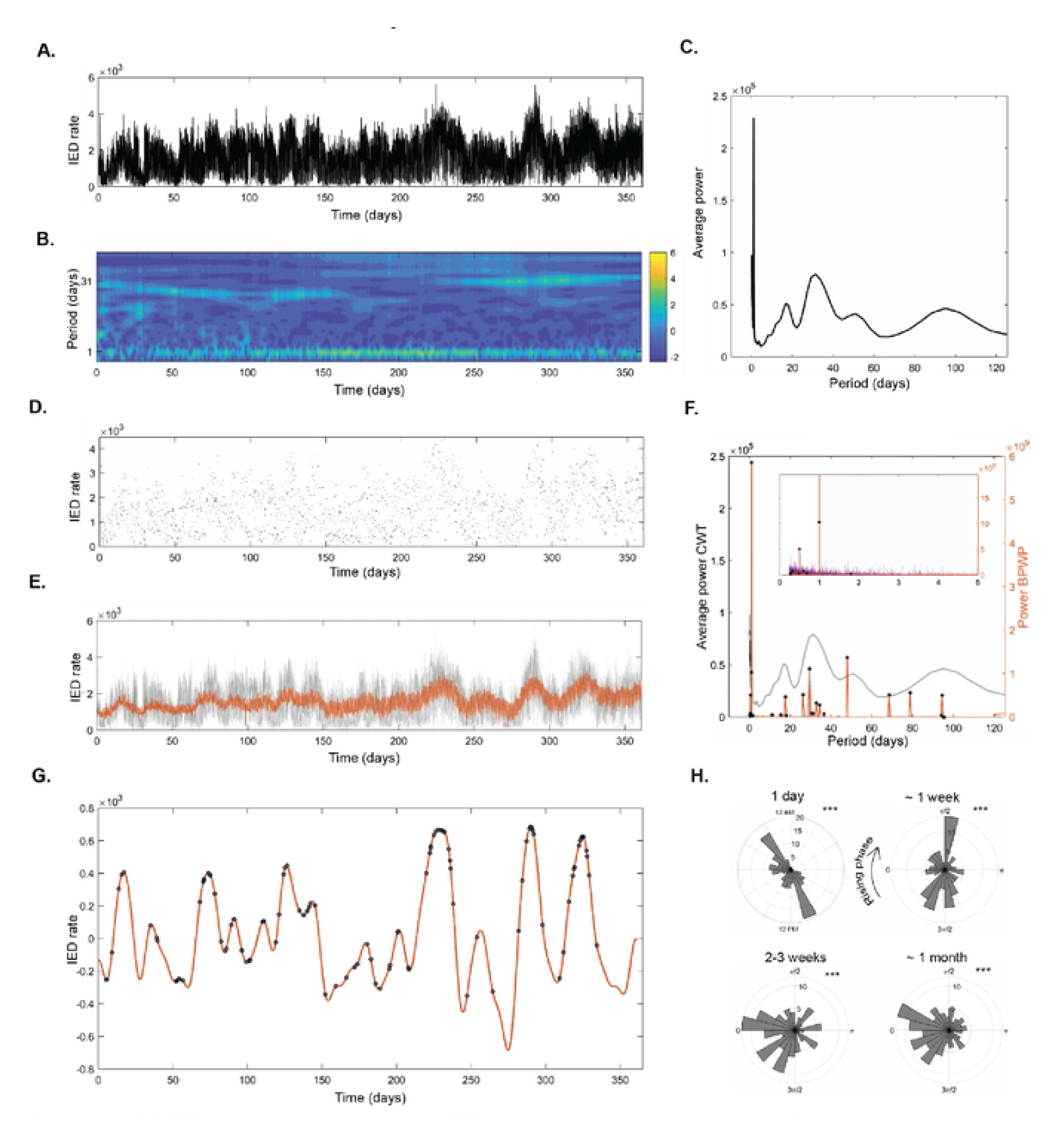
BPWP outputs for real world JED timeseries, participant 1. A) Raw data showing hourly rate of IEDs detected frorn the left hippocarnpus, updated every 20 rninutes. Timeseries consists of over 20,000 sarnples. B) Complex wavelet transfonn (CWT) spectrograrn of the timeseries in A showing power in different cycles. Strong cycles are evident at one day and around on rnonth. C) Average of CWT spectrurn (averaged over tirne frorn the spectrogram in B) shows cycles ofIED rate including periods around one day, two-three weeks, one month, fifty days, and 100 days. D) Randorn samples from the raw data in A averaging at 5 sarnples per day. Timeseries consists of 1,800 total samples. E) Underlying raw data are shown in gray. The method’s estimated reconstruction of the underlying data based on the sparse samples in Dis shown in orange. F) The CWT spectrum for the raw data is shown in gray. The method’s spectral output is shown in orange. Black stars denote significant peaks; peaks whose amplitude was above the 99^th^ percentile of the distribution created by shuffling the input data and re-calculating the method I00 times. The insert shows the spectral outputs from the reshuffling in purple. The amplitude of this noise floor is an order of magnitude smaller than the spectral output from the correctly ordered input data. The narrowband peaks from the method align with the central tendencies of the broadband CWT peaks. G) The method’s estimated reconstruction of the underlying signal using only significant peaks from F with a period longer than two days is in orange. Overlayed black circles denote when seizures occurred. Seizures appear to prefer the peaks of the combined slow cycles derived from the method. H) The method-based reconstruction was filtered in cycle ranges around oneday, one week, two to three weeks, and one month then Hilbert transformed to identify the phase at which seizures occurred for each of these cycles. Polar histograms denoting the phase at which seizures occurred for each of these cycles indicate a cycle-specific phase preference for seizures. Stars denote p < 0.00 I on the Omnibus test for uniformity, indicating that seizure phase is not uniformly distributed.

Figure 4d shows the 1800 random samples (5 per day) from the raw IED rate timeseries of participant 1. The sparse samples represent an over ten-fold decrease in sample density from the standard timeseries. The sparse data were input to BPWP to yield the narrowband DCT power spectrum (Fig. 4f). The BPWP spectrum aligns visually with the main peaks in the CWT spectrum, capturing the highest amplitude circadian and multiday cycles. The “noise floor”, BPWP spectral outputs from repeatedly (100x) reshuffling the original data in time (while preserving the original inter-sample intervals) and re-calculating BPWP, is plotted in purple. Stars denote the spectral peaks whose power exceeded the 99^th^ percentile of the noise floor’s power for that period. The power of the BPWP noise floor (Fig. 4f) is an order of magnitude lower than the BPWP output. The BPWP spectrum and polynomial trend were used to create the signal reconstruction in Figure 4e. This estimated, underlying signal, closely aligns with the original raw data (Fig. 4a) including multiday cycles and the gradual increase in average IED rate over time.

Next, we determined if the reconstructed signal recapitulated the established relationship between cycles of IED rate and seizure occurrence[1]. To visualize the relationship between seizures and the multiday cycles identified by BPWP, we plotted a reconstruction consisting of only the oscillations that were 1) significant and 2) longer than two days (Fig. 4g). For participant 1, the seizures occur near the peaks. Figure 4h shows polar histograms of the phase at which seizures occurred in multiday cycles that are strongly conserved between patients with focal epilepsy[2]. Seizures show cycle-specific phrase preferences (Fig. 4h). The seizure-phase histograms for each cycle were significant on the Omnibus/Hodges-Ajne test[25] (p < 0.001) indicating non-uniform phase distribution and cycle-specific phase preferences. This indicates that the IED cycles identified by BPWP are pathophysiologically relevant.

BPWP outputs for the other two participants are available in Supplementary figures 3 and 4. Overall BPWP performance is similar for participants 1 and 2 (Fig. 4, Supplementary fig. 3), both of which show multiday cycles evident in the raw data. Multiday cycles were less obvious in the raw data for participant 3 and similarly in the BPWP outputs (Supplementary fig. 4) Seizures in all three participants show a clear unimodal or bimodal phase preference within the circadian and multiday cycles.

### Real-world data loss: Impact of sampling density

Examples of different random sampling rates and associated BPWP outputs are shown in Figure 5. For all but the shortest CWT cycles (< 1 day), the scaled offset with the nearest BPWP peak decreases as the number of random samples per day increases, indicating that overall performance improves as more samples are available (Fig. 5h). Offsets stabilize at around 3 samples per day and above. Sampling density results for participants 2 and 3 echo this relationship (Supplementary fig. 5 and 6).

**Figure 5.**
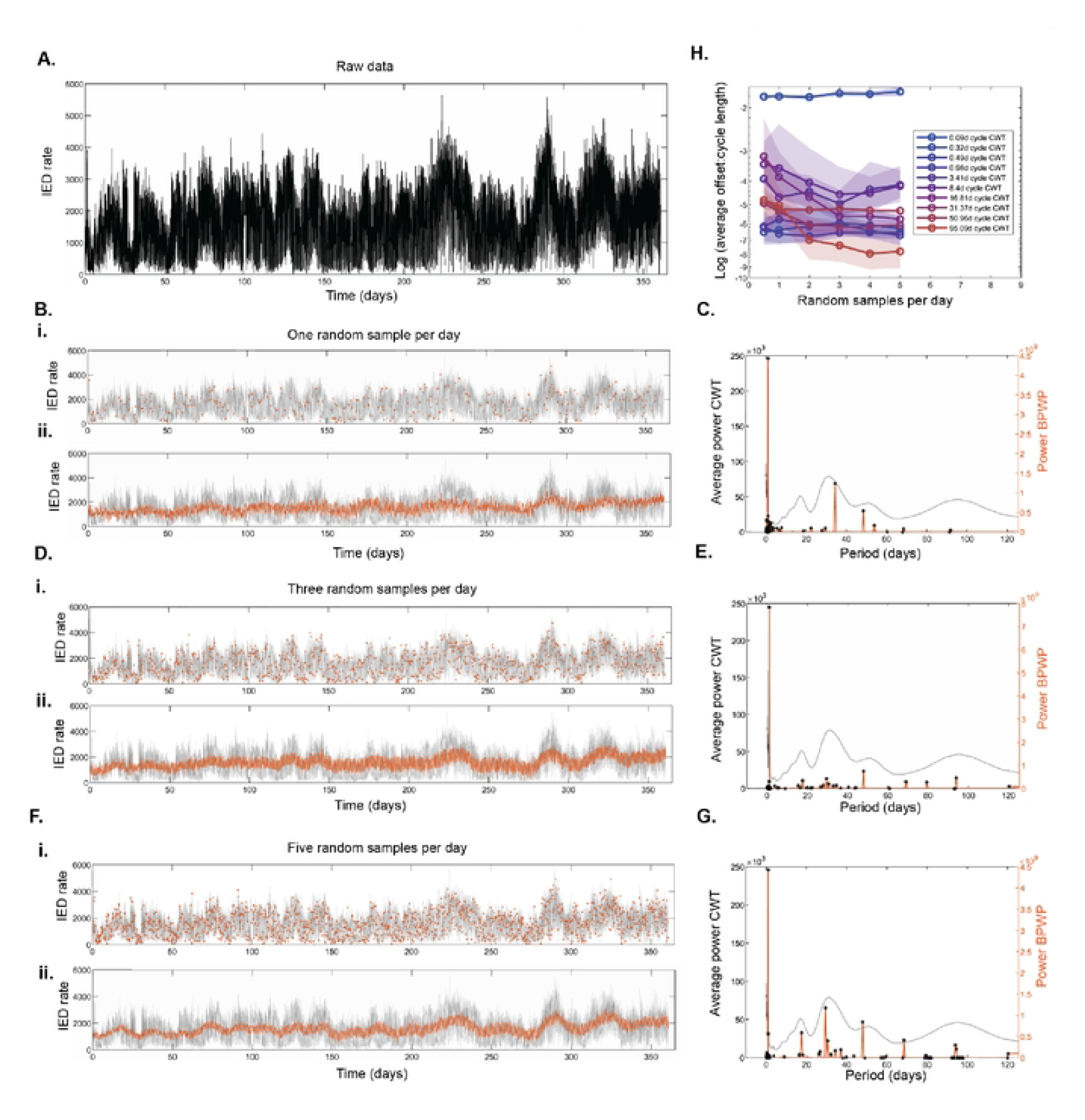
Impact of varying sample density on BPWP outputs, participant 1. A) Raw data showing hourly rate of IEDs detected from the left hippocampus, updated every 20 minutes. Timeseries consists of over 20,000 samples. Bi) Raw data are in gray and one random sample per day is in orange. Bii) Raw data are in gray and the reconstnrcted signal using BPWP outputs based on input data of one randorn sarnple per day is in orange. C) Average complex wavelet transfonn (CWT) spectrum from the raw data in A is in gray. The BPWP spectrum based on one sa1nple per day input is shown in orange. Black stars denote significant peaks; peaks whose an1plitude was above the 99^th^ percentile of the distribution created by shuffling the input data and re-calculating the method 100 times. Randorn samples, signal reconstructions, and BPWP spectra are shown again for sampling rates of three and five per day in D and E and in F and G respectively. Agreernent between BPWP output and raw data and CWT spectra irnproves as the signal is sampled more densely. Part H) shows agreement between the BPWP and CWT spectra as a function of frequency of random sampling. For each sa1npling frequency, the raw dat,avere resarnpled and BPWP was re calculated IO tirnes. For each peak in the CWT spectnun, the offset between the period of the CWT peak and the nearest BPWP peak was calculated in terms of days and divided by the period of the CWT peak. This offset-to-cycle-length ratio was averaged across the IO iterations and plotted as a log value on they axis. The associated frequency of random sampling was plotted on the x axis. Shaded areas denote 95% confidence intervals. The offset ratio decreases and stabilizes as sarnpling density increases.

**Figure 6.**
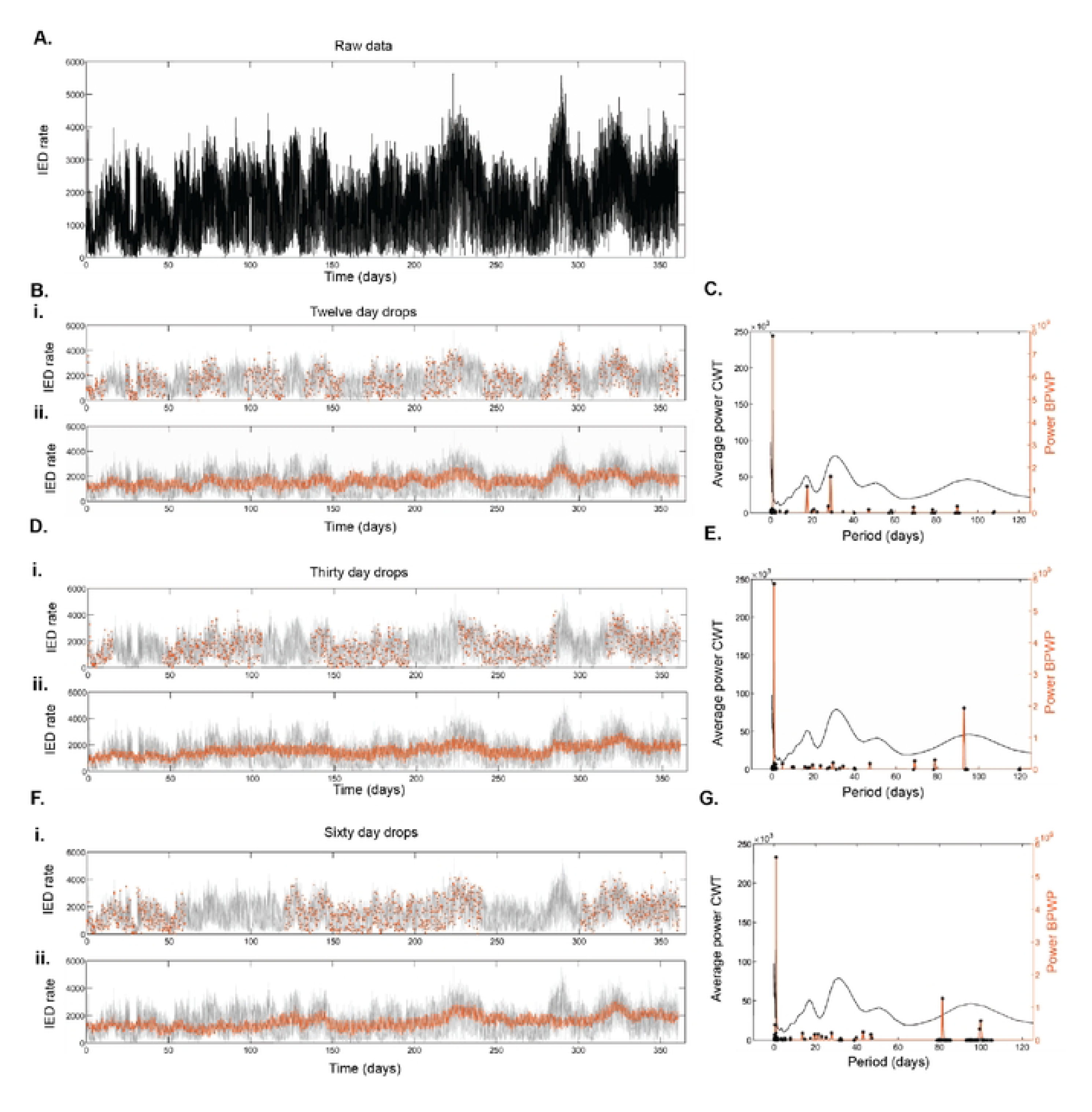
Impact of data drops on BPWP outputs, participant 1. A) Raw data showing hourly rate of IEDs detected from the left hippocampus, updated every 20 minutes. Timeseries consists of over 20,000 samples. Bi) Raw data arein gray and random sampling excluding 12-day data drops are in orange. Bii) Raw data arein gray and the reconstructed signal using model output based on input data with 12-day data drops is in orange. C) Average complex wavelet transform (CWT) spectrum from the raw data in A is in gray. The BPWP spectral output based on the sampling in Bi input is shown in orange. Black stars denote significant peaks; peaks whose amplitude was above the 99th percentile of the distribution created by shuffiing the input data and re-calculating the method I00 times. Data drops of thirty-and sixty-days duration, signal reconstructions, and method spectra are shown in D and E and in F and G respectively. The total number of samples for BPWP is fixed across the conditions at n =I307 which is approximately 4 samples per day assuming no drops.

### Real-world data loss: Impact of data drops

Contiguous data drops of 12, 30, and 60 days duration were created in the IED rate timeseries and the remaining data were sampled at a rate of 5 samples per day. BPWP spectral outputs under these conditions (Fig. 6 c, e, g) varied in peak amplitude and position. BPWP reconstructions during the dropped windows had worse alignment with the original data than for the windows where data were available (Fig. 6 bii, dii, fii). The fidelity of the BPWP reconstructions was better in the case of shorter, more frequent drops (12 days) than for the longer, less frequent drops (60 days). The impact of data drops is shown for participants 2 and 3 in Supplementary figures 7 and 8.

## Discussion

BPWP successfully identified multiday cycles of real-world IED rates from human epileptic hippocampus using a sparsely sampled fraction of the samples required of traditional approaches for frequency decomposition. We anticipate BPWP will enable frequency decomposition of sparse neuro-behavioral timeseries in a variety of applications. BPWP may guide efficient sampling of neural features on implanted devices, extending storage and minimizing patient burden.

Basis pursuit denoising (BPDN)[21, 22], like the similar and at times identical formulation of LASSO[23], leverages the properties of the L1 norm. L1 regularization promotes sparsity and robustness to outliers and conducts built-in feature selection. The degrees of freedom is straightforward to derive as well, and is unbiasedly estimated as the number of nonzero coefficients[26, 27]. Assuming the key assumptions (sparsity and separability), are met, these properties lend themselves to reliable model outputs.

We modified BPDN to include a polynomial representation that manages non-stationarity due to slow changes in signal mean over time. Although low-order polynomial bases have been directly incorporated into sparse recovery approaches before[28, 29], the typical approach to polynomial trends in neural signal processing is to remove them during preprocessing. Polynomial detrending as a preprocessing step is highly debated, as the choice of technique can have a non-negligible influence on the slow fluctuations in the data[30]. By directly including polynomial fitting in the model, we nullify polynomial detrending as a preprocessing step and appropriately consider trends and oscillations in parallel. Non-stationarity in the frequency domain, such as oscillation phase shifts or the emergence and disappearance of oscillations are not explicitly addressed by this model. Our implementation assumes stationarity in the frequency domain.

Future directions include better accommodating these spectral non-stationarities.

There are numerous alternatives to BPDN to detect periodic signals in irregularly sampled data. For example, the Lomb-Scargle periodogram is used widely within astronomy research to identify periodicities in unevenly sampled data[31-33]. Lomb-Scargle and other approaches leverage Fourier transforms and least squares or chi-squared fitting[34], which, compared to L1- regularization can yield less-sparse solutions and increase susceptibility to outliers[31, 32]. Other approaches use continuous-time autoregressive moving average models and Bayesian probability theory[35]. We chose to work from the BPDN framework because it is a convex method and includes L1 regularization which enables conservative estimates of component cycles, but acknowledge that alternatives exist and may be more appropriate for different applications.

The computational efficiency of BPWP is limited by several factors. Large NxN matrices can slow computation times, though their exact dimensions depend on the application and number of coefficients being evaluated. The most time-consuming step of this approach is the preliminary, patient-specific parameter sweep by which the *δ* parameter that constrains the L1 norm is defined. Depending on the machine, parallel processing capabilities, density of the parameter sweep, and N, identifying *δ* can take a day to days. Though, given ongoing rapid advancements in processing capacity, we do not consider this a serious limitation. Nonetheless, the workflow presented here is not currently suitable for real-time computations on clinically implanted devices.

Real-world data loss had differential impacts on the reliability of the BPWP outputs. BPWP spectral outputs were reasonably stable above a sampling frequency of approximately 3 random samples per day for the IED rate data, and remarkably recovered the known underlying circadian and longer timescale cycles in the data. Longer data drops (on the order of months) resulted in variable spectra and inaccurate reconstructions in unsampled epochs. BPWP may be more suitable for circumstances of infrequent but consistent sampling, than for situations with long periods of data loss.

The suitability of BPWP for additional neuro-behavioral applications depends in part on the underlying signal. In simulation, the reliability of cycle detection depended on SNR, sampling rate, and variance. In applications where the ground truth is unavailable, hypothesis driven experiments and cautious interpretation of the model output are required. For example, behavioral timeseries such as Likert scaled ecological momentary assessments from participants who tend to use only a narrow range of the scale (low variance and SNR) are unlikely to reveal cycles. Additionally, the sampling paradigm is an important consideration for retrospective application of BPWP to sparse neuro-behavioral timeseries. BPWP requires irregular samples in time because samples collected at regular intervals, such as a specific time of day, introduce false cycles at the interval duration.

We extrapolated from raw LFP to IED rates derived from LFP, showing that a secondary neuro-behavioral feature can be evaluated with sparse recovery. Comparable sparse recovery approaches in neural signal processing have primarily been applied to signal compression (including LFP), action potentials, and signal transmission[36, 37], as opposed to higher order features such as IED rates. Although sparse recovery systems have been explored for data compression in implantable neural devices[38], our findings support the implementation of feature sampling and storage paradigms with downstream basis pursuit-based reconstruction techniques in mind. Random and sparse sampling of epileptiform features could extend storage capacity in clinically approved devices and enable reliable comparison of electrophysiologically based features with behavioral and multi-domain data.

## Materials and methods

### Application

The purpose of BPWP is to identify oscillations and polynomial trends in sparsely and irregularly sampled neuro-behavioral data. The real-world test data used here are IED rate timeseries. IEDs are defined here as sharp transients detected on LFP recordings from depth electrodes implanted in epileptogenic brain regions (Fig. 1bi)[39].

### Signal

We assume the underlying signal consists of oscillations, a polynomial trend, and gaussian noise (Fig. 2a).

### Approach

The overall approach of BPWP is outlined in Figure 2b. Sparse samples are input to the method which estimates the oscillation coefficients and polynomial coefficients that describe the data. These outputs can then be used to estimate the continuous underlying signal.

### Model

The signal model is described in equations 1 and 2 and Figure 2c,

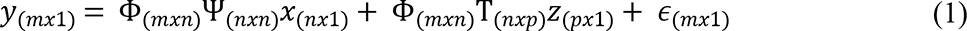

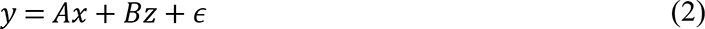

where *y* is a vector of samples of length *m*, Φ is the *m* x *n* binary row subsampling matrix, Ψ is the *n* x *n* discrete cosine transform (DCT) basis, *x* is the length *n* vector of DCT coefficients, Τ is the *n* x p Vandermonde polynomial basis, *z* is the length *p* vector of polynomial coefficients where *p* is equal to the maximum polynomial degree under consideration plus one, and ε is the length m vector of error terms. *A* is the product of Φ and Ψ. *B* is the product of Φ and Τ. Schematic representation of the basis subsampling is depicted in Figure 2dii.

The oscillation basis is a matrix of DCT-II coefficients and determines the frequencies of the component oscillations that can be resolved. The DCT-II expression is shown in equation 3[40]. In equation 3, *s(n)* is the signal at point *n, N* is the total length of the signal *s*, and *δ*_*k*1_ is the Kronecker delta.

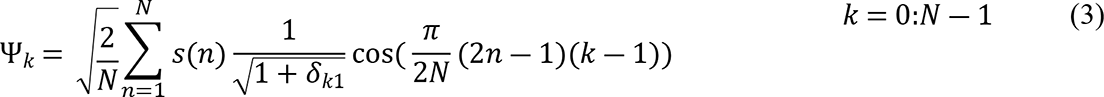

The default frequency representation in the DCT is defined by the sampling rate and the number of points. The associated frequency vector has sparse representation at low frequencies, and dense representation at high frequencies. To ensure that the DCT basis encompasses an adequate density of low frequency coefficients, such as cycles on the order of months or days in length, we incorporated a frequency selection step to the construction of the DCT basis Ψ (equation 4). As in variable density sampling[15], the function *f(k)* can be adapted to optimize the range, density, and central tendency of frequencies evaluated by a transform.

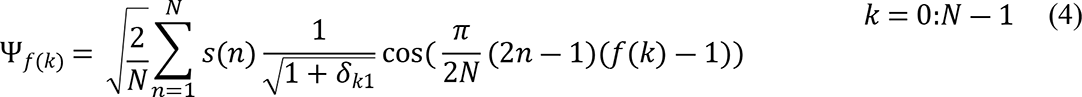

To minimize the disruption of the orthonormal basis structure, we made targeted modifications to the default frequency representation for the DCT via *f(k)*. Briefly, we increased the density of frequency representation within the range of interest on the low frequency end at the expense of randomly removing an equivalent number of frequencies from the high frequency end (Supplementary materials).

### Estimation

BPWP is based on BPDN (equation 5)[21-23]. The parameter *δ* captures error and constrains the L1 norm of the DCT coefficient vector.

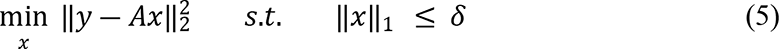

The polynomial representation is incorporated into BPDN in equation 6. This expression has two unknowns, *x* and *z*.

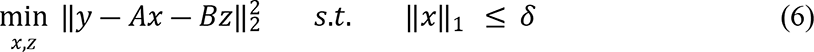

Using variable projection, we reduced the two-dimensional optimization problem to a one-dimensional minimization of *x* (equation 7), then use *x* to solve for *z* (equation 8) [41, 42]. The derivation is available in the supplementary materials.

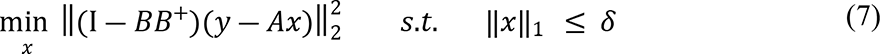

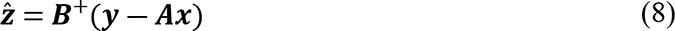

Equations 7 and 8 constitute the core expressions of BPWP. Key assumptions of BPWP are discussed in detail in the supplementary methods.

The DCT and polynomial coefficient outputs can be used to reconstruct the underlying timeseries. Expressions for the signal reconstruction are available in the supplementary materials.

### Features and parameters

Several model components can be adjusted depending on the application. Special considerations are discussed in more detail in the supplementary materials.

### Solver

We used the SDPT3 solver[43, 44] in CVX: Matlab Software for Disciplined Convex Programming (CVX Research Inc.), a system designed for convex optimization in MATLAB (Mathworks®)[45].

### Parameter sweeps

There are many model fits that minimize equation 7. We used patient-specific, 10-fold 75/25 cross validation to identify an appropriate value for *δ* (Supplementary materials). By choosing *δ* based on cross-validation, we select a model that balances over-and under-fitting (the value of *δ* that minimizes mean square error for each participant).

### Determining significance

We repeatedly calculated DCT coefficients *x* by reshuffling the observed samples, *y*, in time (randomly changing y value order with a fixed sampling matrix: preserving the original inter-sample intervals) 100 times to build a nonparametric distribution. Coefficients calculated from the original data were deemed significant if their amplitude was equal to or greater than the 99^th^ percentile of the reshuffling-derived distribution.

### Simulated IED rate timeseries

Simulated data with duration of 12 months and cycle durations comparable to those observed in long-term IED rates[1] were generated. Each sample reflected one hour of simulated IED counts, updated every 20 minutes. Component oscillations included 1, 7, 15, 21, 30, 50, and 100 day periods. Simulated oscillations were present throughout (stationary) the year-long signal duration. A slow trend was included as a first order polynomial.

### Participants

Three, adult, female patients with drug resistant temporal lobe epilepsy were implanted with the investigational Medtronic Summit RC+S™ device[8, 46] as part of an investigational device exemption clinical trial (FDA IDE: G180224, https://clinicaltrials.gov/ct2/show/NCT03946618.) Patients provided written and informed consent in accordance with Institutional Review Board and Food and Drug Administration Requirements.

### IED rate timeseries

One year of continuous IED rate recordings was used from each of the three participants. IEDs were detected from left (the more epileptogenic hemisphere in all participants) hippocampal LFP recordings and logged in one-hour totals updated every 20 minutes[39]. IED rates were collected nearly continuously, with daily hour-long data drops for charging and occasional longer drops due to connectivity issues. For use in the continuous analysis, gaps in the timeseries were interpolated as previously described[1]. The peri-ictal period (two hours prior to seizure, during seizures, and two hours following seizures) was omitted and interpolated as above. For the sparse analysis, the timeseries were randomly sampled according to the relevant paradigm.

### Comparison with complex wavelet transform

Complex wavelet transform (CWT) using Morlet wavelets of the real, original, densely sampled, timeseries was compared with the BPWP spectral outputs (Matlab function cwt, Morlet wavelet, symmetry parameter = 3, time-bandwidth product = 60). The CWT was calculated as described previously[46]. To compare the broadband CWT and narrowband BPWP spectra across sampling conditions, we identified each peak in the CWT spectrum and calculated the offset in terms of period between the CWT peak and the nearest significant BPWP peak detected. The offset was then scaled by dividing it over the length of the CWT cycle. This was repeated for 10 different re-samplings under each condition and averaged.

### Simulating data drops in IED rate timeseries

Contiguous blocks of varying duration (12, 30, and 60 days; 120 total days dropped in each case) were omitted from random sampling to test BPWP performance with data drops. The random daily sampling rate was fixed for the remaining days.

### Seizure phase analysis

Seizures were detected from the raw, continuous LFP recordings[8, 47]. Seizures were defined here as the events whereby seizure detector probabilities exceeded 0.99. To evaluate the phase of seizures within the BPWP reconstruction, the BPWP reconstruction was filtered with a least-squares finite impulse response (FIR) bandpass filter of order (3 * Nyquist frequency) with zero phase shift in filter in cycle ranges that are highly conserved in focal epilepsies[2, 46].: 1, 5-10, 15-25, and 25-35 days. Instantaneous phase was calculated via the Hilbert transform. For each cycle, the phase at which seizures occurred was noted.

### Statistical analysis

The Omnibus (Hodges-Ajne) test for non-uniformity of circular data was used to determine if there was a phase preference for seizures for each multiday cycle evaluated[25].

### Code and processing

Analyses were run on a computer with Intel® Xeon® Silver 4108 CPU @ 1.80 GHz, 188 GB RAM, 16 physical cores, and 32 logical cores, and running Ubuntu version 18.04.6. Code and simulated data are available on GitHub at (https://github.com/irenabalzekas/BPWP). Original clinical data may be made available upon reasonable request and pending compliance with data sharing agreements and appropriate participant privacy protections.

## Acknowledgements

Special thanks to our participants for collecting these data. Additional thanks to Karla Crockett and Cindy Nelson for their vital clinical and administrative efforts. Medtronic provided the Summit RC+S™ system and key scientific collaboration from Abbey Becker PhD, Dave Linde, and Rob Raike PhD, in particular.

## Supporting information

Supplementary methods. (Included separately)

Supplementary figure S1. Subject-specific 10-fold 75/25 cross-validation for delta parameter selection

Supplementary figure S2. Impact of variance, SNR, and cycle length on cycle detection in silico

Supplementary figure S3. BPWP outputs for real world IED timeseries, participant 2

Supplementary figure S4. BPWP outputs for real world IED timeseries, participant 3

Supplementary figure S5. Impact of varying sample density on BPWP outputs, participant 2

Supplementary figure S6. Impact of varying sample density on BPWP outputs, participant 3

Supplementary figure S7. Impact of data drops on BPWP outputs, participant 2

Supplementary figure S8. Impact of data drops on BPWP outputs, participant 3

Supplementary figure S9. Subject-specific 10-fold 90/10 cross-validation for delta parameter selection.

## References

1. Baud MO, Kleen JK, Mirro EA, Andrechak JC, King-Stephens D, Chang EF, et al. Multi-day rhythms modulate seizure risk in epilepsy. Nature Communications. 2018;9(1):88. doi: 10.1038/s41467-017-02577-y.

2. Leguia MG, Andrzejak RG, Rummel C, Fan JM, Mirro EA, Tcheng TK, et al. Seizure Cycles in Focal Epilepsy. JAMA Neurology. 2021;78(4):454–63. doi: 10.1001/jamaneurol.2020.5370.

3. Karoly PJ, Rao VR, Gregg NM, Worrell GA, Bernard C, Cook MJ, et al. Cycles in epilepsy. Nature Reviews Neurology. 2021;17(5):267-84. doi: 10.1038/s41582-021-00464-1.

4. Karafin M, St Louis EK, Zimmerman MB, Sparks JD, Granner MA. Bimodal ultradian seizure periodicity in human mesial temporal lobe epilepsy. Seizure. 2010;19(6):347–51. Epub 2010/06/29. doi: 10.1016/j.seizure.2010.05.005. PubMed PMID: 20580271; PubMed Central PMCID: PMCPMC2921217.

5. Durazzo TS, Spencer SS, Duckrow RB, Novotny EJ, Spencer DD, Zaveri HP. Temporal distributions of seizure occurrence from various epileptogenic regions. Neurology. 2008;70(15):1265–71. doi: 10.1212/01.wnl.0000308938.84918.3f.

6. Herzog AG. Catamenial epilepsy: definition, prevalence pathophysiology and treatment. Seizure. 2008;17(2):151–9. Epub 2008/01/01. doi: 10.1016/j.seizure.2007.11.014. PubMed PMID: 18164632.

7. Griffiths G, Fox JT. RHYTHM IN EPILEPSY. The Lancet. 1938;232(5999):409-16. doi: https://doi.org/10.1016/S0140-6736(00)41614-4.

8. Sladky V, Nejedly P, Mivalt F, Brinkmann BH, Kim I, St. Louis EK, et al. Distributed brain co-processor for tracking spikes, seizures and behavior during electrical brain stimulation. Brain Communications. 2022:fcac115. doi: 10.1093/braincomms/fcac115.

9. Geller EB. Responsive neurostimulation: Review of clinical trials and insights into focal epilepsy. Epilepsy Behav. 2018;88s:11–20. Epub 2018/09/24. doi: 10.1016/j.yebeh.2018.06.042. PubMed PMID: 30243756.

10. Goyal A, Goetz S, Stanslaski S, Oh Y, Rusheen AE, Klassen B, et al. The development of an implantable deep brain stimulation device with simultaneous chronic electrophysiological recording and stimulation in humans. Biosens Bioelectron. 2021;176:112888. Epub 2021/01/05. doi: 10.1016/j.bios.2020.112888. PubMed PMID: 33395569; PubMed Central PMCID: PMCPMC7953342.

11. Kremen V, Brinkmann BH, Kim I, Guragain H, Nasseri M, Magee AL, et al. Integrating Brain Implants With Local and Distributed Computing Devices: A Next Generation Epilepsy Management System. IEEE J Transl Eng Health Med. 2018;6:2500112. Epub 2018/10/13. doi: 10.1109/jtehm.2018.2869398. PubMed PMID: 30310759; PubMed Central PMCID: PMCPMC6170139.

12. Mivalt F, Kremen V, Sladky V, Balzekas I, Nejedly P, Gregg NM, et al. Electrical brain stimulation and continuous behavioral state tracking in ambulatory humans. J Neural Eng. 2022;19(1). Epub 2022/01/18. doi: 10.1088/1741-2552/ac4bfd. PubMed PMID: 35038687; PubMed Central PMCID: PMCPMC9070680.

13. Garudadri H, Baheti PK. Packet loss mitigation for biomedical signals in healthcare telemetry. Annu Int Conf IEEE Eng Med Biol Soc. 2009;2009:2450–3. Epub 2009/12/08. doi: 10.1109/iembs.2009.5333969. PubMed PMID: 19964587.

14. Zhao W, Sun B, Wu T, Yang Z. On-Chip Neural Data Compression Based On Compressed Sensing With Sparse Sensing Matrices. IEEE Transactions on Biomedical Circuits and Systems. 2018;12(1):242–54. doi: 10.1109/TBCAS.2017.2779503.

15. Spielman DM, Pauly JM, Meyer CH. Magnetic resonance fluoroscopy using spirals with variable sampling densities. Magn Reson Med. 1995;34(3):388–94. Epub 1995/09/01. doi: 10.1002/mrm.1910340316. PubMed PMID: 7500878.

16. Bracewell RN. The Fourier transform and its applications: McGraw-Hill New York; 1986.

17. Shumway RH, Stoffer DS, Stoffer DS. Time series analysis and its applications: Springer; 2000.

18. Candes EJ, Romberg JK, Tao T. Stable signal recovery from incomplete and inaccurate measurements. Communications on Pure and Applied Mathematics: A Journal Issued by the Courant Institute of Mathematical Sciences. 2006;59(8):1207–23.

19. Donoho DL. Compressed sensing. IEEE Transactions on Information Theory. 2006;52(4):1289–306. doi: 10.1109/TIT.2006.871582.

20. Ye JC. Compressed sensing MRI: a review from signal processing perspective. BMC Biomedical Engineering. 2019;1(1):8. doi: 10.1186/s42490-019-0006-z.

21. Shaobing C, Donoho D, editors. Basis pursuit. Proceedings of 1994 28th Asilomar Conference on Signals, Systems and Computers; 1994 31 Oct.-2 Nov. 1994.

22. Chen SS, Donoho DL, Saunders MA. Atomic decomposition by basis pursuit. SIAM review. 2001;43(1):129–59.

23. Tibshirani R. Regression Shrinkage and Selection via the Lasso. Journal of the Royal Statistical Society Series B (Methodological). 1996;58(1):267–88.

24. Candès EJ, Wakin MB. An introduction to compressive sampling. IEEE signal processing magazine. 2008;25(2):21–30.

25. Berens P. CircStat: A MATLAB Toolbox for Circular Statistics. Journal of Statistical Software. 2009;31(10):1–21. doi: 10.18637/jss.v031.i10.

26. Zou H, Hastie T, Tibshirani R. On the “degrees of freedom” of the lasso. The Annals of Statistics. 2007;35(5):2173–92, 20.

27. Tibshirani RJ, Taylor J. Degrees of freedom in lasso problems. The Annals of Statistics. 2012;40(2):1198–232, 35.

28. Hernando D, Haldar JP, Sutton BP, Ma J, Kellman P, Liang ZP. Joint estimation of water/fat images and field inhomogeneity map. Magn Reson Med. 2008;59(3):571–80. Epub 2008/02/29. doi: 10.1002/mrm.21522. PubMed PMID: 18306409; PubMed Central PMCID: PMCPMC3538139.

29. Sharma SD, Hu HH, Nayak KS. Accelerated water-fat imaging using restricted subspace field map estimation and compressed sensing. Magn Reson Med. 2012;67(3):650–9. Epub 2011/06/30. doi: 10.1002/mrm.23052. PubMed PMID: 21713983; PubMed Central PMCID: PMCPMC3197950.

30. Woletz M, Hoffmann A, Tik M, Sladky R, Lanzenberger R, Robinson S, et al. Beware detrending: Optimal preprocessing pipeline for low-frequency fluctuation analysis. Hum Brain Mapp. 2019;40(5):1571–82. Epub 2018/11/16. doi: 10.1002/hbm.24468. PubMed PMID: 30430691; PubMed Central PMCID: PMCPMC6587723.

31. Lomb NR. Least-squares frequency analysis of unequally spaced data. Astrophysics and Space Science. 1976;39(2):447–62. doi: 10.1007/BF00648343.

32. Scargle JD. Studies in astronomical time series analysis. II-Statistical aspects of spectral analysis of unevenly spaced data. Astrophysical Journal, Part 1, vol 263, Dec 15, 1982, p 835-853. 1982;263:835–53.

33. VanderPlas JT. Understanding the Lomb–Scargle Periodogram. The Astrophysical Journal Supplement Series. 2018;236(1):16. doi: 10.3847/1538-4365/aab766.

34. Palmer DM. A fast chi-squared technique for period search of irregularly sampled data. The Astrophysical Journal. 2009;695(1):496.

35. Kelly BC, Becker AC, Sobolewska M, Siemiginowska A, Uttley P. FLEXIBLE AND SCALABLE METHODS FOR QUANTIFYING STOCHASTIC VARIABILITY IN THE ERA OF MASSIVE TIME-DOMAIN ASTRONOMICAL DATA SETS. The Astrophysical Journal. 2014;788(1):33. doi: 10.1088/0004-637X/788/1/33.

36. Gurve D, Delisle-Rodriguez D, Bastos-Filho T, Krishnan S. Trends in Compressive Sensing for EEG Signal Processing Applications. Sensors (Basel). 2020;20(13). Epub 2020/07/08. doi: 10.3390/s20133703. PubMed PMID: 32630685; PubMed Central PMCID: PMCPMC7374282.

37. Sun B, Zhao W. Compressed Sensing of Extracellular Neurophysiology Signals: A Review. Front Neurosci. 2021;15. doi: 10.3389/fnins.2021.682063.

38. Suo Y, Zhang J, Xiong T, Chin PS, Etienne-Cummings R, Tran TD. Energy-efficient multi-mode compressed sensing system for implantable neural recordings. IEEE Trans Biomed Circuits Syst. 2014;8(5):648–59. Epub 2014/10/25. doi: 10.1109/tbcas.2014.2359180. PubMed PMID: 25343768.

39. Janca R, Jezdik P, Cmejla R, Tomasek M, Worrell GA, Stead M, et al. Detection of interictal epileptiform discharges using signal envelope distribution modelling: application to epileptic and non-epileptic intracranial recordings. Brain Topogr. 2015;28(1):172–83. Epub 2014/06/28. doi: 10.1007/s10548-014-0379-1. PubMed PMID: 24970691.

40. Ahmed N, Natarajan T, Rao KR. Discrete cosine transform. IEEE Trans Comput. 1974;100(1):90–3.

41. Golub GH, Pereyra V. The Differentiation of Pseudo-Inverses and Nonlinear Least Squares Problems Whose Variables Separate. SIAM Journal on Numerical Analysis. 1973;10(2):413–32.

42. Golub G, Pereyra V. Separable nonlinear least squares: the variable projection method and its applications. Inverse Problems. 2003;19(2):R1–R26. doi: 10.1088/0266-5611/19/2/201.

43. Toh KC, Todd MJ, Tütüncü RH. SDPT3 — A Matlab software package for semidefinite programming, Version 1.3. Optimization Methods and Software. 1999;11(1-4):545-81. doi: 10.1080/10556789908805762.

44. Tütüncü RH, Toh KC, Todd MJ. Solving semidefinite-quadratic-linear programs using SDPT3. Mathematical Programming. 2003;95(2):189–217. doi: 10.1007/s10107-002-0347-5.

45. Boyd MGaS. CVX: Matlab software for disciplined convex programming, version 2.0 beta. http://cvxrcom/cvx. 2013.

46. Gregg NM, Sladky V, Nejedly P, Mivalt F, Kim I, Balzekas I, et al. Thalamic deep brain stimulation modulates cycles of seizure risk in epilepsy. Sci Rep. 2021;11(1):24250. doi: 10.1038/s41598-021-03555-7.

47. Sladky V, Kremen V, McQuown K, Mivalt F, Brinkmann BH, Gompel JV, et al., editors. Integrated human-machine interface for closed-loop stimulation using implanted and wearable devices. 2022 IEEE International Conference on Systems, Man, and Cybernetics (SMC); 2022 9–12 Oct. 2022.

